# A scalable estimator of SNP heritability for Biobank-scale data

**DOI:** 10.1101/294470

**Authors:** Yue Wu, Sriram Sankararaman

## Abstract

**Motivation:** Heritability, the proportion of variation in a trait that can be explained by genetic variation, is an important parameter in efforts to understand the genetic architecture of complex phenotypes as well as in the design and interpretation of genome-wide association studies. Attempts to understand the heritability of complex phenotypes attributable to genome-wide SNP variation data has motivated the analysis of large datasets as well as the development of sophisticated tools to estimate heritability in these datasets.

Linear Mixed Models (LMMs) have emerged as a key tool for heritability estimation where the parameters of the LMMs, i.e., the variance components, are related to the heritability attributable to the SNPs analyzed. Likelihood-based inference in LMMs, however, poses serious computational burdens.

**Results:** We propose a scalable randomized algorithm for estimating variance components in LMMs. Our method is based on a MoM estimator that has a runtime complexity 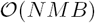 for N individuals and *M* SNPs (where *B* is a parameter that controls the number of random matrix-vector multiplications). Further, by leveraging the structure of the genotype matrix, we can reduce the time complexity to 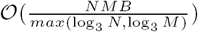.

We demonstrate the scalability and accuracy of our method on simulated as well as on empirical data. On standard hardware, our method computes heritability on a dataset of 500, 000 individuals and 100, 000 SNPs in 38 minutes.

**Availability:** The RHE-reg software is made freely available to the research community at: https://github.com/sriramlab/RHE-reg

## 1 Introduction

A central question in biology is to understand how much of the variation in a trait (phenotype) can be explained by genetics as opposed to environmental factors. The heritability of a trait is a central notion in quantifying the contribution of genetics to the variation in a trait. The heritability of a trait refers to the proportion of variation in the trait that can be explained by genetic variation (Visscher *et al*., 2008). The narrow-sense heritability (*h*^2^) refers to the proportion of trait variation that can be explained by a linear function of genetic variation (Almasy and Blangero, 1998). Beyond understanding the genetic basis of a phenotype, heritability determines the power of genetic association studies to detect genetic variants associated with a phenotype, the accuracy of using genetic data to predict phenotypes, as well as the response of a phenotype to natural and artificial selection (Houle, 1992).

While family-based studies enabled the estimation of heritability of a wide variety of traits, the availability of genome-wide genetic variation data has enabled a direct estimation of the heritability associated with genotyped SNPs, termed SNP *heritability*. Initial attempts to estimate heritability from genomic data focused on the variation in a trait the could be explained by SNPs that were discovered to be significantly associated with the trait in a genome-wide association study (GWAS). These estimates were found to severely under-estimate the narrow-sense heritability, a phenomenon known as *missing heritability.* A major insight into the mystery of missing heritability emerged in Yang *et al.* (2010a) who showed that using all genotyped SNPs jointly to explain variation in a trait led to a substantially larger estimate of heritability than from SNPs that were found to be associated in GWAS. Subsequent analyses suggest that much of missing heritability could be explained by the presence of a large number of SNPs of weak effects that has, in turn, motivated analyses of larger datasets.

Linear Mixed Models (LMMs) has emerged as a key analytically technique for estimating the heritability of complex traits using genome-wide SNP variation data. Beyond their application in estimating SNP heritability, LMMs are widely used in association tests where they are used to control for population stratification (Kang *et al.,* 2008a; Yu *et al.,* 2006; Lippert *et al.,* 2011; Loh *et al.,* 2015b; Zhou and Stephens, 2014), in phenotype and disease risk prediction (Zhou *et al.,* 2013; Makowsky *et al.,* 2011; Wray *et al.,* 2013; Speed *et al.,* 2012; Yang *et al.,* 2010a), and in understanding the relative contribution of genomic regions to variation in a trait of interest (Yang *et al.,* 2010a; Makowsky *et al.,* 2011; Wray *et al.,* 2013). A key step in the application of LMMs is the estimation of their parameters, *i.e.,* often referred to as variance components. Estimation of variance components is a computationally challenging problem on genomic datasets containing large numbers of individuals and SNPs. The most commonly used method for variance components estimation in LMMs relies on maximizing the likelihood of the parameters. Often, a related estimator, known as the restricted maximum likelihood (REML) estimator, is preferred due to a reduced bias relative to maximum likelihood estimators. Both maximum likelihood as well as REML estimation, however, rely on computationally intensive optimization problems. While a number of methods have been proposed to improve the computational efficiency of REML estimators (Yang *et al.,* 2010b; Kang *et al.,* 2008b; Lippert *et al.,* 2011; Loh *et al.,* 2015b,a; Pirinen *et al.,* 2013), all of these methods rely on iterative optimization algorithms that do not scale well to Biobank-scale datasets consisting of millions of individuals genotyped at tens of millions of SNPs. Further, REML has been shown to yield biased estimates of heritability in ascertained case-control studies (Golan *et al.,* 2014; Chen, 2014).

## 1.1 Our contributions

We propose a scalable randomized algorithm to estimate variance components of a linear mixed model. Our method is based on Haseman-Elston (HE) regression (Haseman and Elston, 1972; Elston *et al.,* 2000; Chen *et al.,* 2004; Bulik-Sullivan, 2015), a Method-of-Moment (MoM) estimator of the heritability of a phenotype. The HE regression estimator, like other MoM estimators, tends to be statistically less efficient compared to REML. On the other hand, HE regression is computationally attractive as it leads to a set of linear equations in the variance components that can be solved analytically. While this property of HE regression is appealing, a key computational bottleneck in the application of HE regression is the computation of an *N* × *N* matrix that summarizes the relationship between all *N* pairs of individuals in the dataset. As a result, the computation and memory requirements of HE scale quadratically with the number of individuals.

Our randomized HE regression (RHE-reg) estimator relies on the observation that the key bottleneck in HE regression can be replaced by multiplying the *N* × *M* (individuals × SNPs) matrix of genotypes with a small number, *B,* of random vectors. This leads to a randomized estimator with runtime 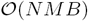 and memory requirements 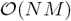. Further, we leverage the observation that the genotype matrix has entries in a finite set, *i.e.,* {0, 1, 2}so that the time complexity of matrix-vector multiplication reduces to 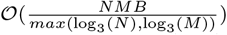 (Liberty and Zucker, 2009). This additional gain in efficiency can be substantial when the number of SNPs or individuals is large. For example, in the UK Biobank, *N* is of the order of 10^5^ while *M* is of the order of 10^6^. Thus, we propose an estimator of variance components with runtime 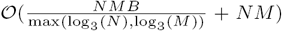 and memory requirement 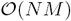.

We apply the RHE-reg estimator to the problem of estimating SNP heritability. We show that our method yields unbiased SNP heritability estimates. While our method is statistically inefficient compared to REML (both because it is moment-based as well as the added randomization), we show in practice that the statistical inefficiency is minimal, particularly for large sample sizes. Further, our method is substantially more computationally efficient so that it can be effectively applied to whole-genome genotype data from hundreds of thousands of individuals. REML has been shown to yield biased estimates of heritability in ascertained case-control studies (Chen *et al.,* 2004; Golan *et al.,* 2014) while the RHE-reg estimator can also be applied in this setting.

Finally, since variance component analysis is of interest beyond heritability estimation, the RHE-reg estimator can enable rapid estimation of variance components in all of the settings in which LMMs are used.

## 2 Method

We observe genotypes from *N* individuals at *M* SNPs. The genotype vector for individual *i* is a length *M* vector denoted by *g*_*i*_ ∈ {0, 1, 2}^*M*^. The *j*^*th*^ entry of *g*_*i*_ denotes the number of minor allele carried by individual *i* at SNP *j*. Let *G* be the *N* × *M* genotype matrix where 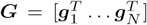. ***X*** is a *N* × *M* matrix of standardized genotypes obtained by centering and scaling each column of ***G*** so that 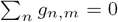 and 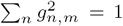 for all *n* ∈ {1,…, *N*}. Let *y* is an *N*-vector of phenotypes and *β* be an *M*-vector of SNP effect sizes.

### 2.1 Linear Mixed Model

We assume the vector of phenotypes *y* is related to the genotypes by a linear mixed model:

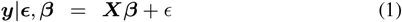

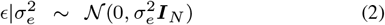

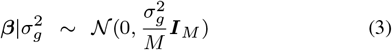

Here *y* is centered so that 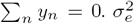 is the residual variance while 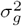 is the variance component corresponding to the *M* SNPs. The SNP heritability is defined as 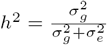.

In this model, we have 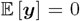 while the population covariance of the phenotype vector *y* is:

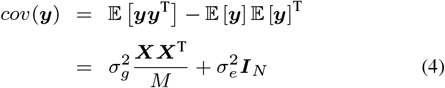

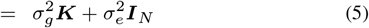

Here 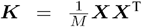 is the genetic relatedness matrix (GRM) computed from all SNPs. One approach to estimate the SNP heritability is Haseman-Elston regression (Haseman and Elston, 1972) which is a MoM estimator obtained by equating the population covariance to the empirical covariance (several variants of Haseman-Elston regression have been proposed; what we consider here is HE-CP (Sham and Purcell, 2001)). The empirical covariance of the phenotype vector *y* is estimated by *yy*^T^.

The MoM estimator is obtained by solving the following ordinary least squares problem (see Appendix A.1 for details):

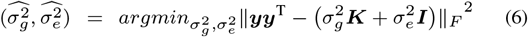

The MoM estimator satisfies the normal equations:

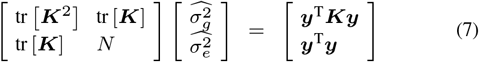

Solving the normal equations requires computing 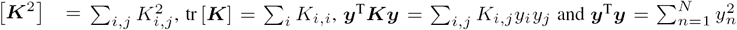. The GRM ***K*** can be computed in time 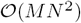 and requires 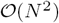 memory. Given the GRM, computing each of the coefficients for the normal equation requires 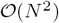 time. Finally, given each of the coefficients, we can solve analytically solve for the 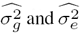. Indeed, we can write

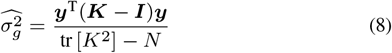

Thus, the key bottleneck in solving the HE regression lies in computing the GRM.

### 2.2 RHE-reg: A randomized estimator of heritability

Given that 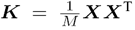, we can compute the quantities tr 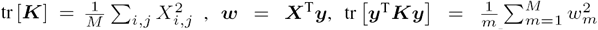. For standardized genotypes, tr [***K***] = *N* while tr 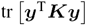 can be computed in 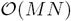 time.

The one remaining quantity that we need to compute efficiently is tr [*K*^2^]. Given a *N* × *N* matrix ***A*** and a random vector *z* with mean zero and covariance *I*_*N*_, we use the following identity to construct a randomized estimator of the trace of matrix ***A*** (see Appendix A.2 for a proof):

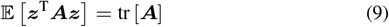

Equation 9 leads to the following unbiased estimator of the trace of ***K***^2^ given *B* random vectors, *z*_1_,…, *z*_*B*_, drawn independently from a distribution with zero mean and identity covariance matrix ***I***_*N*_:

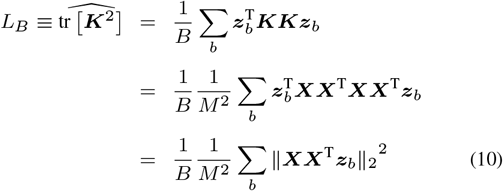

In practice, we draw each entry of *z* independently from a standard normal distribution. We note that the estimator *L*_*B*_ involves two matrix-vector multiplications of *N* × *M* matrix repeated *B* times for a total runtime of 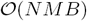.

The RHE-reg estimator 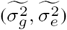 is obtained by solving the Normal equations (Equation 7) by replacing tr [*K*^2^] with *L*_*B*_.

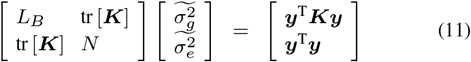

The RHE-reg estimator of the SNP heritability is then obtained by 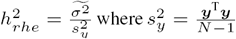 is the unbiased estimator of the phenotypic variance.

### 2.3 Sub-linear computations

The key bottleneck in the RHE-reg is the computation of *L*_*B*_ which involves repeated multiplication of the normalized genotype matrix ***X*** by a real-valued vector. Leveraging the fact that each element of the genotype matrix ***G*** takes values in the set {0, 1,2}, we can improve the complexity of these multiplication operations from 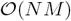 to 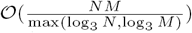 using the Mailman Algorithm (Liberty and Zucker, 2009).

#### 2.3.1 The Mailman algorithm

Consider a *M × N* matrix ***A***^*T*^ whose entries take values in {0,1, 2}. Assume that the number of SNPs *M =* log_3_(*N*). The naive way to compute the product ***A***^*T*^ *b* for any real-valued vector *b* takes *O*(*log*_3_(*N*)* *N*) time.

The mailman algorithm decomposes the matrix ***A*** as ***A***^*T*^ = *U*_*n*_*P*. *U*_*n*_ is a *log*_3_(*N*) × *N* matrix whose column contains all possible vectors over {0, 1, 2} of length *log*_*3*_(*N*). And *P* is an indicator matrix, where entry *P*_*i,j*_ = 1 if the *i*^*th*^ column is the same as *j*^*th*^ column in matrix 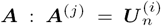. The decomposition of matrix ***A*** takes 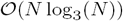 time. The desired product ***A***^*T*^ *b* is computed in two steps as *c* = ***Pb*** followed by *U*_*n*_*c*, each of which can be computed in only 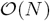 operations (Liberty and Zucker, 2009).

For a matrix ***A***^*T*^ with *M* > [log_3_(*N*)], we partition ***A***^*T*^ into 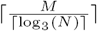 sub-matriceseachofsize 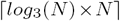 each of which can be multiplied in time 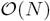 for a total computational cost of 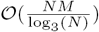.

#### 2.3.2 Application of the Mailman algorithm to RHE-reg

Now consider the standardized genotype ***X***, which could be written as *X =* (***G*** — M)∑, where ***M*** is a matrix where the *i*^*th*^ column contains the sample mean of the *i*^*th*^ 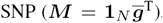, and ∑ is an *M × M* diagonal matrix, with the inverse of variance of each SNP as the diagonal entries.

Thus, when we compute 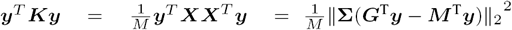 in Equation 14, computing 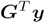 using the Mailman algorithm takes 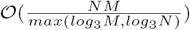 operations. Similarly, to compute each term in the sum of the randomized estimator of tr [***K***^2^] (Equation 10), ***X***^*T*^ *z*_*b*_, we can substitute *X*^*T*^ *z*_*b*_ with ∑*G*^*T*^ *z*_*b*_ — *∑M*^*T*^ *z*_*b*_. The first term *∑G*^*T*^ *z*_*b*_ can again be computed using 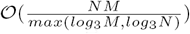 using the Mailman algorithm, and the second term *∑M*^*T*^ *z*_*b*_ is equivalent to scaling the *N*-vector *z*_*b*_ which can be computed in time 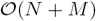.

### 2.4 Computing the Standard Error

We show in Appendix A.4 that the variance of the RHE-reg estimator of 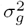 can be approximated by the variance of the exact HE-regression estimator with an additional contribution due to the randomization:

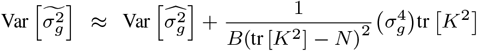

Here *B* is the number of samples used and *z* is a random vector with mean zero and identity covariance matrix. For samples with low-levels of relatedness, we can assume ***K*** *≈* ***I*** and our estimates of 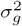 and tr [***K***^2^] to estimate the variance. Further, we show in Appendix A.4 that we can estimate the variance (and hence, the standard error) of the RHE-reg estimator in sub-linear time without assuming that ***K*** *≈**I**.*

### 2.5 Some remarks on the RHE-reg estimator

1. The RHE-reg is biased as we show in Appendix A3 with a bias that decreases with *B.* In practice, the bias appears to be small (see Figure 1).
2. Equation 3 assumes an infinitesimal model for the phenotype. However, all our results only dependonthe second momentofthe SNP effect sizes. Thus, the RHE-reg estimator can yield valid estimates for non-infinitesimal architectures.
3. In a number of settings, it is desirable to include covariates, such as age or sex, in the analysis. This changes the model in Equation 3 to:

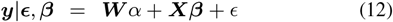

Here ***W*** is a *N × C* matrix of covariates while α is a C-vector of coefficients. In this setting, we transform Equation 12 by multiplying by the projection matrix 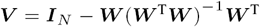:

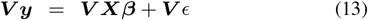

The RHE regression estimator applied to Equation 13 then must satisfy the following moment conditions:

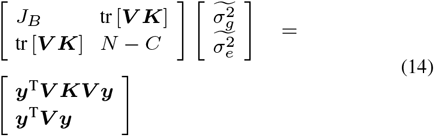

Here *J_B_* is a randomized estimator of tr [***V K V K***] analogous to Equation 10. The cost of computing the RHE-reg estimator now includes the cost of computing the inverse of ***WW***^T^ as well as multiplying ***W*** by a real-valued vector for an added computational cost of 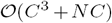. Typically, the number of covariates *C* is small (tens to hundreds) so that the presence of covariance does not greatly increase the computational burden.
4. The variance components model (Equation 3 and 5) can be extended in a straightforward manner to more than two variance components. A number of recent studies have explored the utility of these models to partition heritability based on functional annotations as well as other categories.
5. The accuracy and the runtime of RHE-reg depends on the choice of the number of random vectors *B.* In practice, we find that the estimator is highly accurate with a small *B ≈* 100 even for moderate sample sizes *N* ≈ 5,000 as we show empirically (Figure 3). Further, for larger sample sizes, even smaller values of *B* should be adequate. It is also possible to choose increasing values of *B* and to terminate when the estimate of tr [***K***^*2*^] does not change considerably. We have not explored this option in detail in this work.

**Fig. 1.**
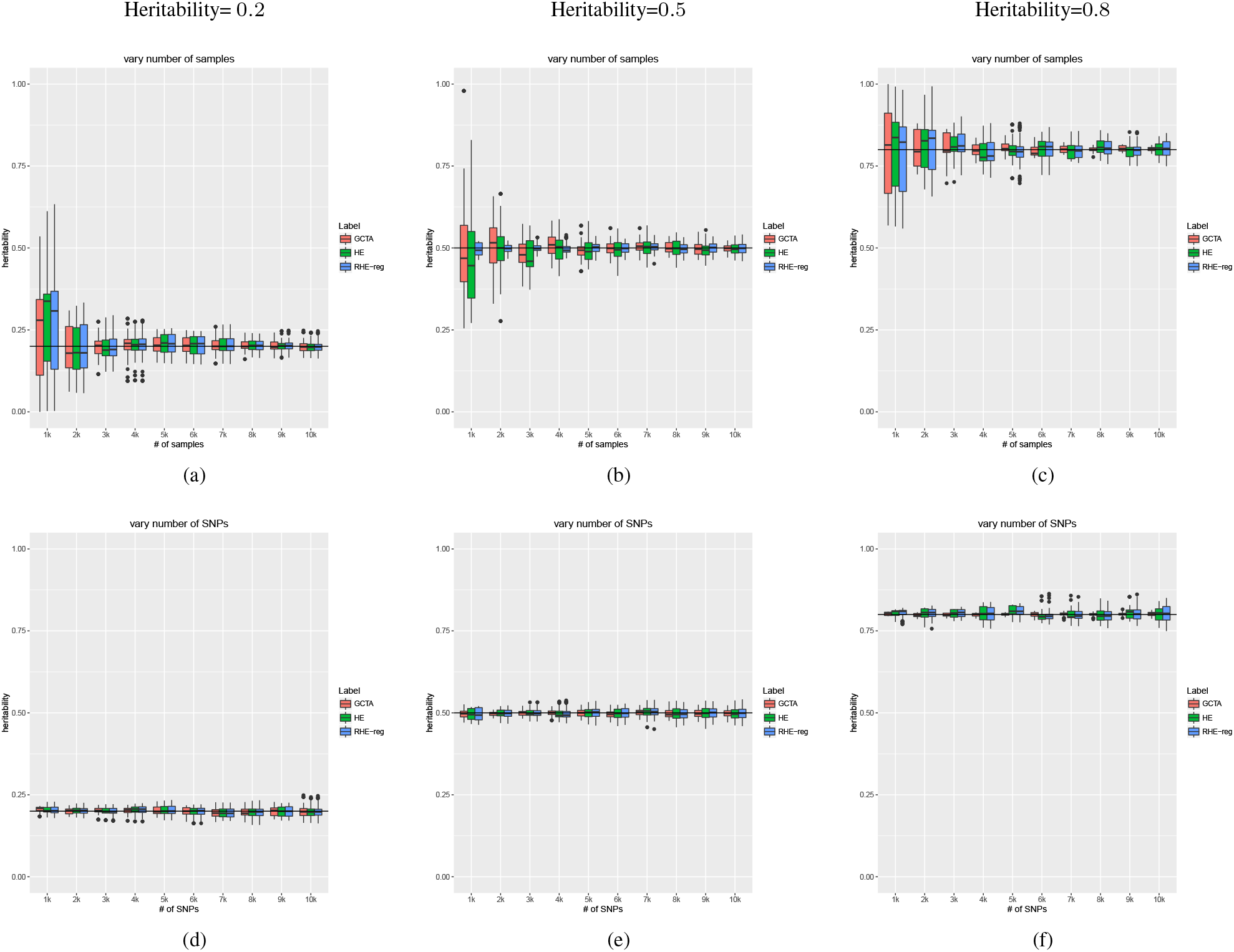
RHE-reg accurately estimates heritability: In the first series of Figures 1(a)-(c), we simulated genotypes with varying sample size while fixing the number of SNPs to 10, 000. The phenotype in each of the Figures 1(a),1(b) and 1(c) is simulated with true heritability of 0.2, 0.5, and 0.8 respectively. The second series of figure 1(d)-1(f) considers genotype data with varying number of SNPs while the number of samples is fixed at 10, 000. All three methods that we evaluated (GCTA, HE-reg and RHE-reg) have similar accuracies. GCTA which estimates the REML has smaller standard errors when the heritability is large (*h*^2^ = 0.80). For lower values of true heritability (*h*^2^ = 0.20, *h*^2^ = 0.50), the estimates from REML, HE-regression and RHE-reg are comparable. HE and RHE-reg have similar variance suggesting that randomization only makes a minor contribution to the statistical accuracy.

## 3 Results

### 3.1 Simulations

We performed simulations to measure the performance of RHE-reg to other methods for heritability estimation in terms of accuracy, running time and memory usage. We compared RHE-reg to two methods for computing REML estimates: GCTA (Yang *et al*., 2010b) (which implements an exact numerical optimization algorithm to compute the REML) as well as implementations of Haseman-Elston regression.

### 3.2 Accuracy

In our first set of simulations, we compared the accuracy of RHE-reg to our implementation of exact HE-regression as well as GCTA, an implementation that computes the REML. We simulated genotypes assuming each SNP is drawn independently from a Binomial distribution with allele frequency that is sampled uniformly from the interval (0, 1). Given the genotypes, we simulated phenotypes under an infinitesimal model, *i.e.*, with effect size at each SNP drawn independently from a normal distribution with mean zero and variance equal to the heritability divided by the number of SNPs. We considered different values for the true SNP heritability of the phenotype to be 0.2, 0.5, and 0.8.

In our first series of experiments, we fixed the number of SNPs at *M* = 10, 000 and varied the number of individuals *N* = 1*k*, 2*k*…10*k*. In the second series of experiments, we varied the number of SNPs *M* = 1*k*, 2*k*…10*k* while fixing the number of individuals to be *N* = 10, 000. We repeated each experiment 100 times in order to assess the variance of each of the estimators. We estimated heritability using RHE-reg with *B* = 100 random vectors.

Figure 1 compares the estimates of each of the three methods (RHE-reg, HE-regression, and GCTA) to the true heritability. First, we observe that all three methods obtain estimates of heritability that are quite close to each other as well as to the true heritability across the range of parameters explored. Second, RHE-reg and HE-regression are virtually indistinguishable in the variance of their estimates in each configuration. This suggests that the randomization makes a negligible contribution to the statistical accuracy of the MoM estimators. In some cases, RHE-reg even has a smaller variance than HE-regression. Thirdly, as expected, REML obtains estimators that are closer to the true heritability compared to either of the MoM estimators for a high value of true heritability. For lower values of true heritability (*h*^2^ = 0.20, *h*^2^ = 0.50), the estimates from REML, HE-regression, and RHE-reg are comparable. This result is also expected given that REML is asymptotically equivalent to MoM when the phenotypic correlation between individuals is small (Sham *et al*., 2000; Sham and Purcell, 2001). Finally, the sample size has a bigger effect than the number of SNPs on the accuracy of each of the methods, consistent with theory (Visscher *et al*., 2014).

### 3.3 Computational Efficiency

In the second set of simulations, we compared the runtime and memory usage of different methods. We compared RHE-reg to two REML methods, GCTA (Yang *et al*., 2010b) and BOLT-REML (Loh *et al*., 2015a) (a computationally efficient approximate method to compute the REML) as well as an exact MoM method MMHE (Ge *et al*., 2017). In this experiment, we simulated genotype data consisting of 100, 000 SNPs over sample sizes of *N* = 10*k*, 20*k*, 30*k*, 50*k*, 100*k* and 500*k* and then simulated phenotypes corresponding to the genotype data. For each dataset, we ran RHE-reg with *B* = 100 random vectors. We performed all comparisons on an Intel(R) Xeon(R) CPU 2.10GHz server with 128 GB RAM. All computations were restricted to a single core, capped to a maximum runtime of 12 hours and a maximum memory of 128 GB.

Figure 2 shows that both GCTA and MMHE do not scale to large sample sizes due to the requirement of computing and operating on a genetic relatedness matrix (GRM) that scales quadratically with *N*. GCTA could not complete its computation when running on *N* = 100K individuals while MMHE did not complete its computation on *N* = 50*K*. BOLT-REML and RHE-reg scale linearly with sample size. However, RHE-reg is an order of magnitude faster than BOLT-REML. For example, on a dataset of a size of 500*K* individuals, RHE-reg computed the heritability in about 30 minutes compared to 400 minutes for BOLT-REML. Figure 2 shows that RHE-reg is memory efficient as well.

**Fig. 2.**
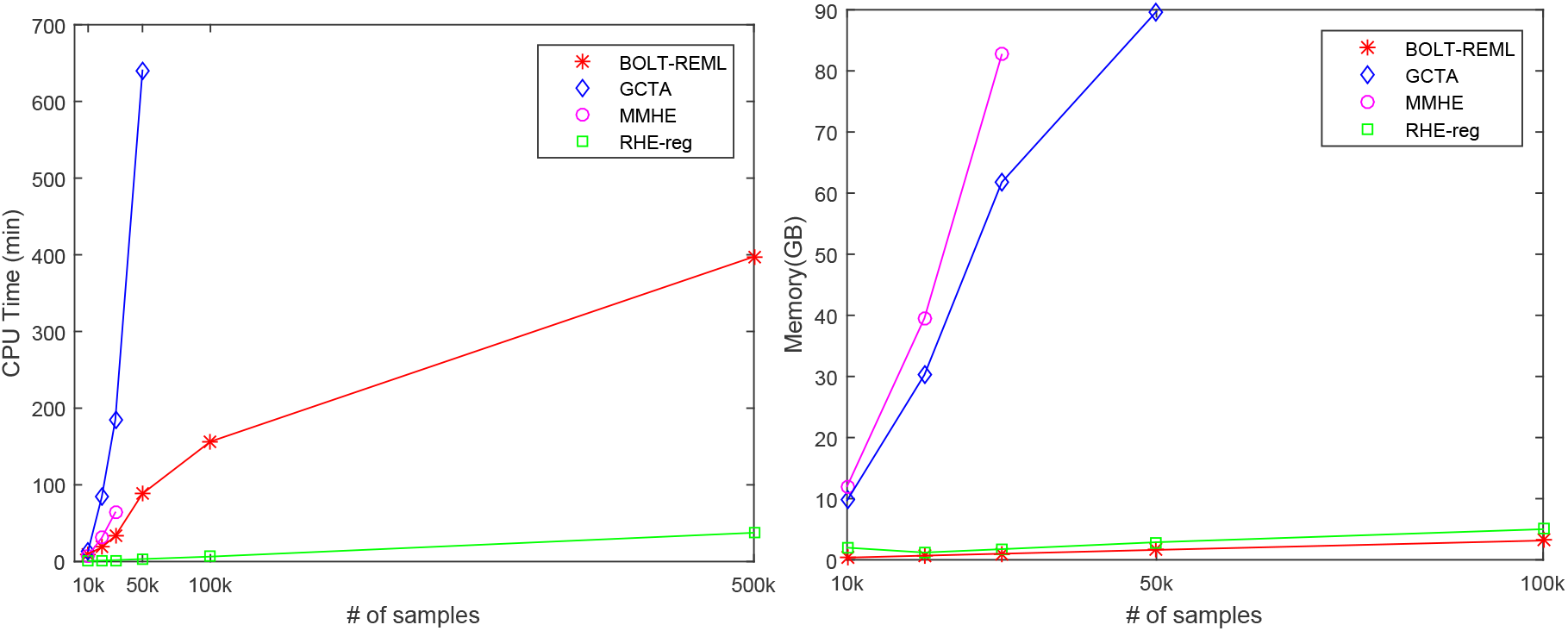
RHE-reg is efficient: We measured the run time and memory usage of methods for heritability estimation as a function of the number of samples while fixing the number of SNPs to 100, 000. We performed all comparisons on an Intel(R) Xeon(R) CPU 2.10GHz server with 128 GB RAM. All computations were restricted to a single core, capped to a maximum runtime of 12 hours and a maximum memory of 128 GB. In Figure 2(a), GCTA could not finish computation on 100K samples. For MMHE, the computation stopped at sample size of 50k due to memory constraints. Although BOLT-REML scales linearly, RHE-reg is significantly faster. In Figure 2(b), we observe RHE-reg and BOLT-REML have scalable memory requirements.

**Fig. 3.**
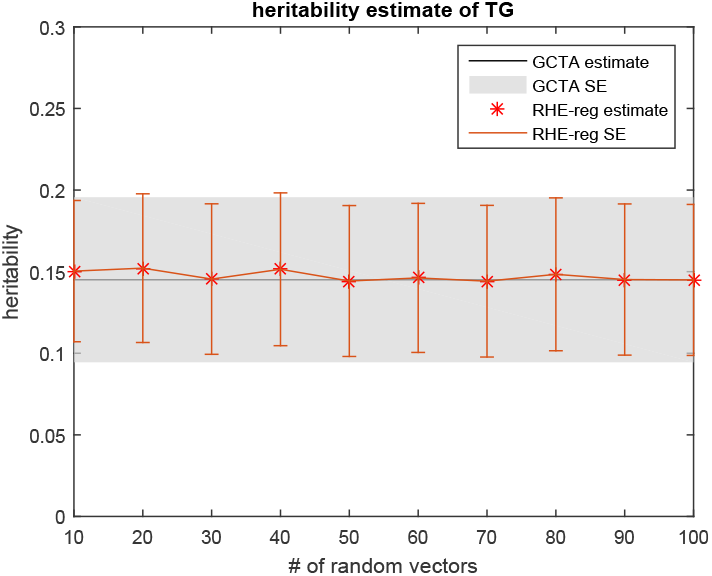
Impact of the number of random vectors on the accuracy of RHE-reg: We ran RHE-reg with a different number of random vectors *B*, and compared the point estimate and standard error to GCTA. The gray area indicates the standard error computed by GCTA. As RHE-reg use more random vectors, the estimate converges. In fact, even with 10 random vectors, the point estimation is accurate.

### 3.4 Application to real data

We compared the statistical accuracy and run time of BOLT-REML, GCTA, and RHE-reg on the Northern Finland Birth Cohort (NFBC) dataset. The NFBC dataset contains 315, 529 SNPs and 5, 326 individuals after applying standard filters (minor allele frequency > 0.05 and Hardy-Weinberg Equilibrium p-value < 0.01) (Sabatti *et al*., 2009). We applied these methods to estimate the heritability of three phenotypes that were assayed in this dataset: triglycerides (TG), high-density lipoprotein (HDL) and body mass index (BMI).

We compared the runtime, point estimates of the heritability as well as standard errors for each of the three methods. We computed RHE-reg with *B* = 100 random vectors. As shown in the Table 1, the heritability estimates of RHE-reg are concordant with the other methods while being an orderofmagnitude fastertocompute. Wenote that the NFBC dataset has a sample size *N* ≈ 5,000 so that we expect RHE-reg to be more accurate on larger datasets. The standard error estimates can also be computed in sub-linear time (see Appendix A.4).

**Table 1.** The estimates ofheritability from RHE-reg are consistent with those from GCTA and BOLT-REML on the NFBC data while RHE-reg is substantially faster. We estimate the heritability of phenotypes such as triglycerides (TG), high-density lipoprotein (HDL), and body mass index (BMI) in the NFBC data set.

|  | Method |  |  |  |  |  |
| --- | --- | --- | --- | --- | --- | --- |
|  | GCTA |  | BOLT-REML |  | RHE-reg |  |
| | Runtime | $h_g^2$ | Runtime | $h_g^2$ | Runtime | $h_g^2$ |
|  | (min) | (se) | (min) | (se) | (min) | (se) |
| TG | 11.28 | 0.145<br>(0.051) | 8.87 | 0.148<br>(0.051) | 1.61 | 0.145<br>(0.052) |
| HDL | 10.81 | 0.325<br>(0.051) | 9.72 | 0.326<br>(0.051) | 1.30 | 0.349<br>(0.052) |
| BMI | 10.85 | 0.237<br>(0.051) | 9.29 | 0.235<br>(0.051) | 1.29 | 0.200<br>(0.052) |

### 3.5 Understanding the computational efficiency of RHE-reg

Our implementation of RHE-reg relies on two ideas to obtain computational efficiency: i) the use of a randomized estimator of the trace, and ii) the Mailman algorithm for fast matrix-vector multiplication. To explore the contribution of each of these ideas, we compared the runtimes of a MoM estimator with no randomization (HE-reg), RHE-reg using standard matrix-vector multiplication and RHE-reg using the Mailman algorithm. Table 2 shows the runtimes of each of these variants on the NFBC data. We see that the biggest runtime gain arises from applying the randomized estimator (faster by a factor of 10-12 relative to HE-reg) while the application of the Mailman algorithm reduces the runtime further by a factor of 2 (Table 1).

**Table 2.** We compare the run time for HE-reg as well as the run time for RHE-reg that does not rely on the Mailman algorithm. The major gain in computational efficiency arises from the application of the randomized trace estimate.

|  | Runtime | No Mailman | No randomized<br>trace estimate |
| --- | --- | --- | --- |
|  | (min) | (min) | (min) |
| TG | 1.61 | 3.70 | 38.5 |
| HDL | 1.30 | 2.60 | 36.2 |
| BMI | 1.29 | 2.68 | 36.7 |

### 3.6 Accuracy of RHE-reg as a function of the number of random vectors B

To explore the impact of the choice of the number of random vectors *B* on the accuracy of RHE-reg, we compared the heritability estimates of RHE-reg to those obtained from GCTA for the triglyceride (TG) phenotype as a function of *B*. We find good concordance between the estimates from RHE-reg and GCTA even for values of *B* as low as 10 suggesting that RHE-reg could be even faster in practice with little loss in accuracy 3.

## 4 Discussion

We proposed a scalable estimator of heritability which is a randomized version of the Haseman-Elston regression (RHE-reg) estimator. The RHE-reg estimator is based on performing a small number of multiplications of the genotype matrix with random vectors with mean zero and identity covariance. Using the properties of the genotype matrix, we can compute this estimator using the Mailman algorithm in 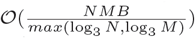 time on a dataset containing *N* individuals, *M* SNPs and with a small number of *B* random vectors. We show that this estimator achieves similar accuracy as REML-based methods on both simulated and real data. RHE-reg can be effectively applied to whole-genome genotype data of hundreds of thousands of individuals for rapid variance components estimation. Furthermore, RHE-reg is an unbiased estimator and thus can also be applied to ascertained case-control studies.

## Acknowledgments

We thankXiangZhouand the reviewers fortheir valuable feedback. SSwas supported in part by is supported in part by NIH grants R00GM111744, R35GM125055, NSF Grant III-1705121, an Alfred P. Sloan Research Fellowship, and a gift from the Okawa Foundation.

## Appendix

### A.1 Method of Moments

The method of moments principle obtains estimates of the model parameters such that the theoretical moments match the sample moments. In our model, the first theoretical moment, 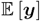, is 0 by definition while the corresponding sample moment is also zero since we standardized the phenotypes. The second sample momentis *yy*^*T*^ and the second theoretical moment is 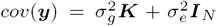. Thus, the method-of-moments (MoM) estimator of 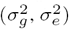 is obtained by searching for values of 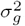, 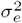 such that the sample and theoretical moments are close, *i.e.,* by solving an ordinary least squares (OLS) problem:

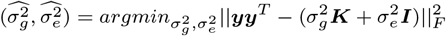

Since the Frobenius norm of a matrix 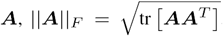, the OLS problem can be re-written as:

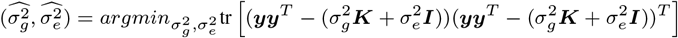

which leads to Equation 6.

### A.2 Randomized Estimator of trace of a Matrix

For a *N × N* matrix, ***A***, a randomized estimator of tr 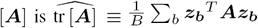, where *z*_*b*_ are i.i.d. random vectors with each entry drawn from a standard normal distribution. To see this:

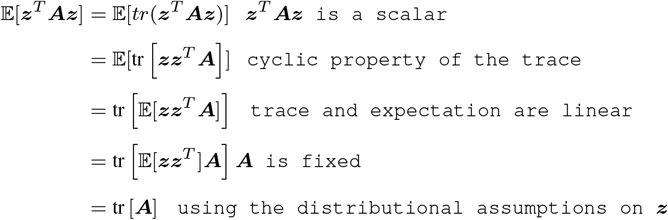

### A.3 Bias of the RHE-reg Estimator

Our estimator of tr 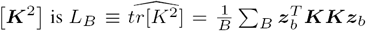. The RHE-reg estimators for 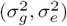 are given by: 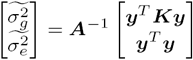 where 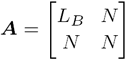.

We first compute the expectation of this estimator:

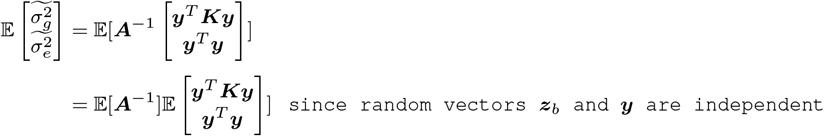

We know that 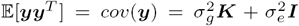. We can compute 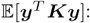

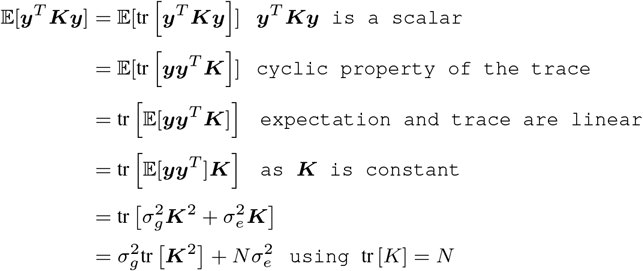

And for 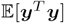, we have;

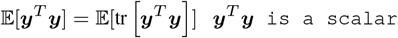

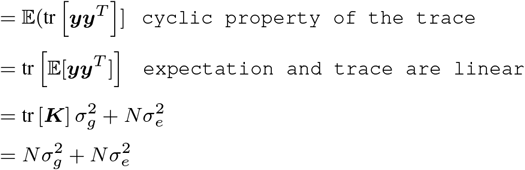

Defining 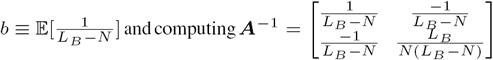, we have

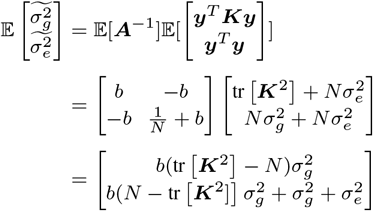

We approximate 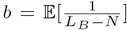 using Taylor expansion. As we have: 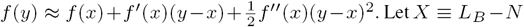, and thus 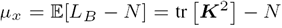. We have 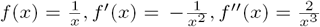.

Thus:

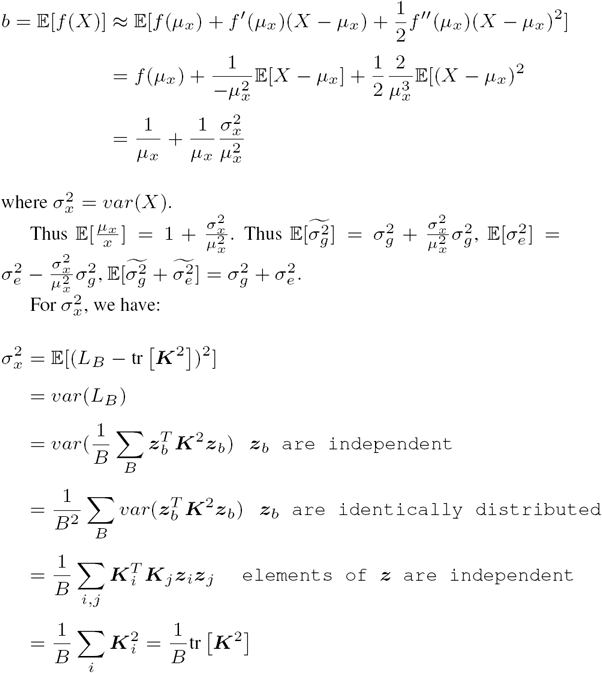

Here *K*_*i*_ is the *i*^*th*^ column of K.

Thus, substituting 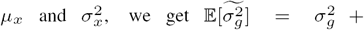 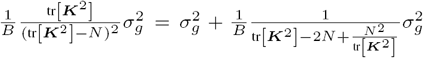. The bais of the estimator decreases with larger number of random vectors *B*.

### A.4 Standard Error Estimate for the RHE-reg estimator

We define 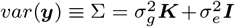. As we know 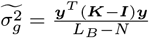. Let 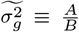 where 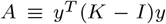 and *B* ≡ *L*_*B*_ − *N*. Define 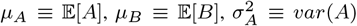 and 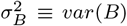. From Lemma 2, we have

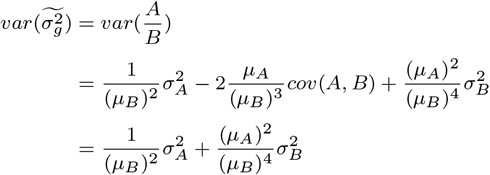

as *A, B* are independent. By using Lemma 1, we have:

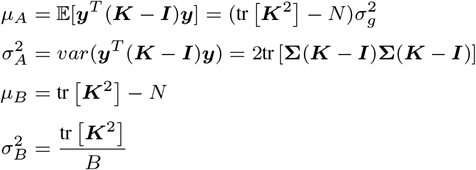

Thus we have:

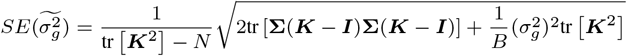

In order to estimate the standard error of 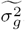 we use the plug-in estimator:

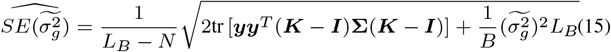

Each term in this estimator could be efficiently computed in 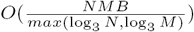.

### A.5 Useful identities

Lemma 1: For a random vector *z* that is distributed according to a multivariate normal distribution: 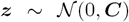 and for symmetric matrices *A* and *B.*

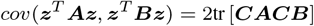

Thus

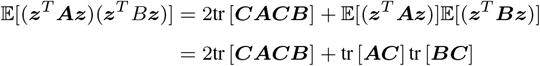

Lemma 2: For two random variables, *A* and *B,* where B is either discrete or has support 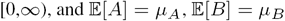.

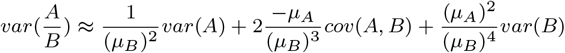

